# Revisiting the Fungal ITS2 Sequencing Data from the Human Microbiome Project

**DOI:** 10.1101/2025.07.01.662555

**Authors:** Xinyu Wang, Wenhao Zhou, Wanglong Gou, Ju-Sheng Zheng

**Affiliations:** Affiliated Hangzhou First People’s Hospital, School of Medicine, Westlake University, Hangzhou, China; Zhejiang Key Laboratory of Multi-Omics in Infection and Immunity, School of Medicine, Westlake University, Hangzhou, China; Westlake Laboratory of Life Sciences and Biomedicine, Hangzhou, China

## Abstract

While the Human Microbiome Project (HMP) significantly advanced our understanding of the human microbiome, our revisiting of its fecal ITS2 sequencing data reveals two significant issues. First, technical variations in sequencing or data processing introduce batch effects in downstream analyses. Second, we discovered a mismatch between the reported and actual amplification primers, which requires careful consideration when interpreting, reusing, or referencing HMP ITS2 data and methodology.

## Main

The Human Microbiome Project (HMP) has significantly advanced our understanding of the human microbiome in two key ways: generating extensive reusable, high-quality data and helping standardize analytic pipelines of human microbiome research^1–3^. Beyond bacteria, HMP extended its research to viral and fungal communities^4,5^, enhancing our understanding of these minority species in the gut and other body sites. Notably, in 2017, Nash et al. published a study characterizing fungal communities in HMP samples using ITS2, 18S rRNA gene amplicon sequencing, and metagenomic sequencing—work that has profoundly influenced subsequent population-based studies of gut fungi (nearly 900 citations)^5^. However, our recent analysis of the HMP ITS2 raw data revealed two concerning issues. First, the presence of two different sequencing methods—or unreported reads trimming—introduced batch effects in downstream analyses. Second, using our newly developed pipelines, we discovered a discrepancy between the reported and actual amplification primers. These issues warrant careful consideration when analyzing the related data or referencing the associated methodology.

We analyzed paired-end sequencing files labeled as ITS2 sequencing from 324 samples (PRJNA356769, NCBI). Our initial analysis revealed that while most samples had read lengths of 300 bases (consistent with the paper’s reported paired-end 300, PE300), 25 samples (7.7%) had read lengths of 250 bases, indicating either trimming or paired-end 250 (PE250) sequencing (Supplementary Table 1). For clarity, we refer to these two types as PE300 and PE250 in the following analysis. Upon examining the sequencing depth of these two types of samples, we found that PE250 samples had significantly higher sequencing depth than PE300, even after removing 14 PE300 samples that contained fewer than 1,000 reads (Figure 1a, P = 0.0018). This further suggests that these two sample types likely came from different sequencing batches.

**Fig. 1:**
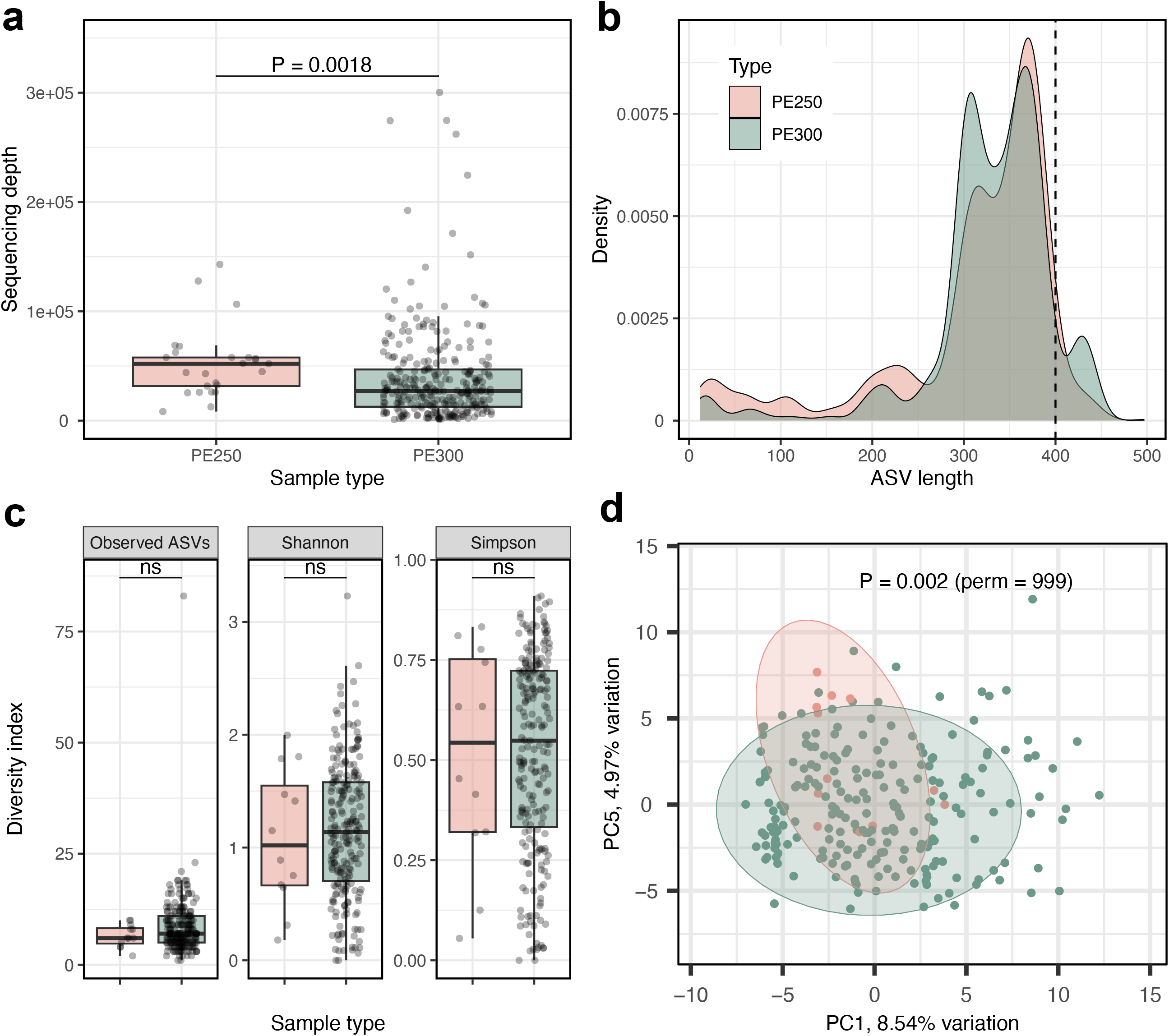
Technical variations in HMP ITS2 sequencing introduce batch effects in downstream analyses. **a**, Sequencing depth (reads per fastq file) of PE250 (n = 25) and PE300 (n = 285) samples containing more than 1,000 reads. P-value were determined using the Kruskal-Wallis test. **b**, Distribution of ASV lengths from the DADA2 pipeline comparing PE250 (n = 25) and PE300 (n = 292) samples, with 7 samples excluded from annotation due to low sequencing depth. **c**, Alpha diversity (Observed ASVs, Shannon index, and Simpson index) of non-mushroom fungal ASVs from PE250 (n = 12) and PE300 (n = 223) samples, each containing more than 5,000 non-mushroom fungal reads. P-values were determined by Kruskal-Wallis test. ns, P > 0.05. **d**, Beta diversity (Aitchison distance) between fungal communities from PE250 (n = 12) and PE300 (n = 223) samples, each containing more than 5,000 non-mushroom fungal reads. P-value was determined using PERMANOVA with 999 permutations. PC, principal component.

Since ITS2 region length varies considerably (50-1000 bp), shorter read lengths could fail to capture longer ITS2 sequences, affecting downstream analysis^6,7^. Further analysis revealed that PE250 samples had a maximum ITS2 Amplicon Sequence Variant (ASV) length of 443, compared to 497 for PE300, with PE300 samples containing more ASVs exceeding 400bp (Figure 1b). Additionally, though fungal alpha diversity remained similar between sample types (Figure 1c), their community structures differed significantly (Figure 1d, P = 0.002, permutations = 999). These findings indicate that HMP ITS2 raw data likely came from different sequencing strategy or underwent inconsistent processing, potentially compromising downstream analyses.

The high length variation in fungal ITS2 regions normally causes linked-adapter issues on reads when sequencing cycles exceed insert sequence length. Processing ITS amplicon data requires removing adapter sequences to prevent artificial sequences from affecting denoising, chimera identification, and taxonomic annotation^7,6^. Using primer sequences as adapter identification sites is standard practice, as adapter sequences typically flank amplification primers. However, using the mainstream adapter detection tool Cutadapt^8^, we were surprised to find that almost no reported ITS2 primer (ITS3: GCATCGATGAAGAACGCAGC and ITS4: TCCTCCGCTTATTGATATGC) could be detected on either forward or reverse reads, even though we confirmed most samples were untrimmed (all reads within the same sample were either 300bp or 250bp). This finding contrasted sharply with other dataset (Guangzhou Nutrition and Health Study, GNHS^9^) that used identical primers and sequencing strategy, where fewer than 1% of reads lacked primer sequences. In another well-defined fecal ITS1 sequencing dataset (MK-SpikeSeq^10^), we successfully detected the reported PCR primer sequences in 99.8% of forward reads and 75.5% of reverse reads. We hypothesized that our inability to detect primer sequences in the HMP data was due to poor sequencing quality. However, even after relaxing adapter identification thresholds, over 99.9% of reads of HMP still failed to detect this primer pair. Moreover, quality filtering (method) retained most files (634/648) and reads (24,291,384/24,767,324), indicating that the absence of expected ITS2 amplification primer sequences was not due to poor sequencing quality (Supplementary Table 2).

Suspecting incorrect primer reporting of HMP ITS2 dataset, we sought to identify the actual primers used for amplification. Since there were no tools available for de novo primer identification, we developed Amplicon Primer Hunter (APHunter, Method). This tool identifies consistent consecutive sequences within high quality reads of each fastq file, as adapter barcodes and PCR primers should be uniform within files. After identifying consistent sequences (red area in plot), the tool performs a BLAST search against a primer database to identify potential primers in the reads (Figure 2a). Additionally, we built a comprehensive database containing widely used fungal ITS high-throughput sequencing primers (n = 78, including ITS3 and ITS4, Supplementary Table 3) through literature review^7,11,12^. We validated APHunter’s performance using GNHS and MK-SpikeSeq datasets. In the GNHS dataset, ITS3 and ITS4 primers were detected in all 1,631 R1 and R2 files, with only two files showing a single mismatch (Supplementary Table 4 and 5). In the MK-SpikeSeq dataset, all R1 files contained the ITS1F primer with zero mismatches. Additionally, 251 out of 308 R2 files showed significant BLAST results, all containing ITS2 (a reverse primer for ITS1) with zero mismatches (Supplementary Table 6 and 7).

**Fig. 2:**
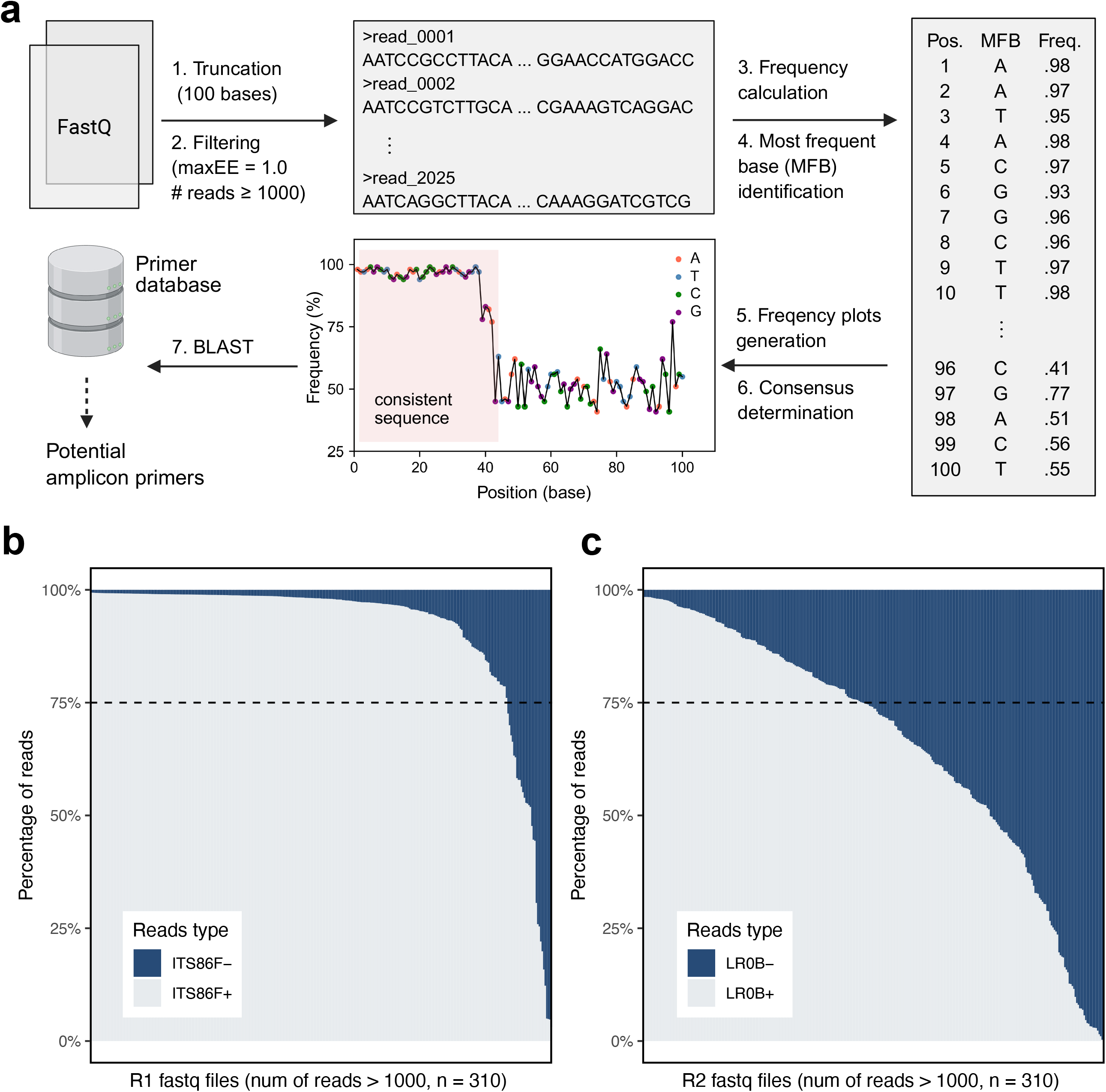
Primer identification reveals mismatch between reported and actual primers in HMP ITS2 data. **a**, Schematic diagram of the APHunter analysis pipeline. Figure created with BioRender.com. **b, c**, Percentage of reads containing ITS86F or LR0B primers in each R1 and R2 fastq file. ITS86F+ represents reads where the ITS86F sequence was detected, while ITS86F-represents reads where it was not detected. The same notation applies to LR0B+ and LR0B-.

Using APHunter, we analyzed primers in the HMP ITS data. Of the 310 R1 files that passed filtering, 296 contained consistent sequences longer than 25bp. BLAST analysis identified 238 sequences with significant hits (E < 0.001), but none matched ITS3. Instead, all best hits consistently matched ITS86F, with most showing 100% sequence similarity (Supplementary Table 8). Similarly, for R2 files, 308 passed filtering, with 278 containing consistent sequences longer than 25bp. Of these, 240 showed significant hits (none matching ITS4), with 237 matching LR0B, though many had mismatches (140 hits ≥1, Supplementary Table 9). Further Cutadapt analysis using relaxed parameters confirmed that more than 90% of R1 files (>1,000 raw reads) contained the ITS86F sequence in at least 75% of their reads (ITS86F+, Figure 2b). This indicates that ITS86F, not ITS3, was likely the forward primer used for amplification.

However, only 44.7% of R2 files (>1,000 raw reads) had more than 75% of reads containing the LR0B sequence (LR0B+, Figure 2c), suggesting the authors used either alternative primers not present in our library or a combination of different primers.

Next, we developed an alternative annotation strategy to circumvent primer-dependent adapter removal while addressing the linker-adapter issue (Method). The analysis yielded 1,510 ASVs from 10,658,025 reads (86.1% of raw reads), with 1,381 ASVs containing ITS2 regions (82.3% of raw reads). BLAST searches against the ITS primer database revealed no significant ITS3 or ITS4 hits (E < 0.01, Supplementary Table 10). However, ITS86F was present in 86.7% of ASVs containing ITS2 regions (88.6% of ASV table reads) with minimal mismatches (≤2), while only 32.6% of ASVs showed significant hits for LR0B (65.4% of ASV table reads). These findings further confirmed that the HMP study likely did not use ITS3 and ITS4 as their ITS2 amplification primers.

As a cornerstone of human microbiome research, HMP’s work has influenced countless subsequent studies. Over the past eight years, research groups have extensively studied the human gut mycobiome using large-scale ITS sequencing^13,14^. Our findings reveal that the mycobiome study of HMP likely misreported which ITS2 region amplification primers were used. This discrepancy is significant since primer choice critically affects high-throughput sequencing results^6,7,15^. Ironically, ITS3 and ITS4 primers have become widely adopted, despite not being optimal for ITS2 region sequencing^6,7^.

In conclusion, our findings suggest that HMP’s fecal ITS2 sequencing data should be interpreted with caution. The combination of PE300 and PE250 sequencing methods or potential undocumented sample processing introduced significant batch effects. Moreover, the mismatch between reported and detected primers requires careful consideration when citing HMP’s methodology or making comparisons across studies that use different primers.

## Method

### Adapter detection

Cutadapt (v5.0) was used to detect primer sequences. For paired-end data, we used the linked adapter option with two parameter sets: a default set (maximum error rate = 0.1, minimum overlap = 0.9) and a relaxed set (maximum error rate = 0.2, minimum overlap = 0.75). For R1 files, we set the forward primer ITS3 (GCATCGATGAAGAACGCAGC) as adapter 1 (required and not anchored) and the reverse complement sequence of ITS4 (GCATATCAATAAGCGGAGGA) as adapter 2 (optional and not anchored). For R2 files, ITS4 and the reverse complement sequence of ITS3 were set as adapter 1 and 2 with the same dapter search parameters, respectively. We discarded untrimmed reads (--discard-untrimmed) and empty reads (--minimum-length 1) and used SeqKit to calculate the number of reads and bases before and after processing. For single-end data, the regular 5’ adapter option (-g) was used with default parameters set.

### APHunter development

APHunter is a Python-based tool that identifies primers in amplicon sequencing data from fastq files. The workflow follows these steps: First, SeqKit truncates input reads to the first 100 bases. Next, VSEARCH filters out low-quality reads using a maximum expected error (maxEE) threshold of 1.0. The tool then evaluates read counts per sample, discarding samples with fewer than 1,000 reads to ensure reliable primer identification. For remaining samples, it calculates nucleotide frequencies (A, T, C, G) at each base position across truncated reads. A frequency plot shows the most frequent base (MFB) at each position, visualizing base distribution across read length. The tool determines consensus sequences using MFBs and truncates them when five consecutive MFBs show frequencies below 0.75—indicating less conserved regions. These consensus sequences are compiled into a.fasta file and compared against a primer database (e.g., ITS primer database) using BLASTn-short. Significant matches identify potential amplification primers.

APHunter is available through conda (https://anaconda.org/westraingroup/aphunter) and its source code is hosted on GitHub (https://github.com/WeStrainGroup/APHunter).

### ITS2 sequencing data annotation

Raw paired-end fastq files were filtered and trimmed using the filterAndTrim function in the DADA2 R package with default parameters, except maxEE = 5, to remove reads with excessive errors. The DADA2 algorithm then inferred ASVs from R1 and R2 files. Denoised forward and reverse reads were merged using the MergePairs function. We set trimOverhang = TRUE to trim the overhanging sequences that flank the alignment region of forward and reverse reads— necessary because some amplicons were shorter than the read length (300 or 250 bp)—as an alternative to the Cutadapt linked adapter option. The resulting structure of each ASV contained barcode-forward primer-5.8S-ITS2-28S-reverse primer-barcode. Since the barcode and flanking 5.8S and 28S rRNA genes show no or poor taxonomic resolution and affect chimera identification, we used ITSx to extract the ITS2 region from ASVs and reconstructed the ASV table before chimera filtering and taxonomic annotation. ASVs that failed to show ITS2 features were discarded from further analysis, as these likely originated from nonspecific amplification. The widely used naive Bayesian classifier plug in the DADA2 package (assignTaxonomy) was used for taxonomy assignment. We used the UNITE general FASTA release for eukaryotes (dynamic version of 19.02.2025) as our reference database to identify fungi, plants, and other eukaryotic sequences simultaneously. This comprehensive approach is necessary for ITS sequencing since common primers typically co-amplify plant DNA sequences, particularly from dietary sources. Since fungal genomes typically contain multiple distinct ITS copies, which can inflate richness estimates as DADA2 identifies these intragenomic ITS2 sequence variants, we clustered the ASVs at 98.5% similarity using VSEARCH and reconstructed the ASV table (ref).

The phyloseq object was constructed for subsequent filtering and analysis. ASVs with non-fungal kingdom-level annotations or unclear phylum-level annotations were filtered out, and counts were set to zero for ASVs with relative abundance below 1/10,000 in each sample. After removing mushroom reads from the phyloseq object using a customized script, only samples containing more than 5,000 non-mushroom fungal reads were retained for subsequent analysis.

### Alpha and beta diversity analysis

The estimate_richness function from the phyloseq R package was used to calculate alpha diversity indices (Observed, Shannon, and Simpson) based on the filtered ASV table. For beta diversity analysis, we aggregated the ASV table to genus level and performed centered log-ratio transformation (CLR, pseudocount = 1) using the decostand function of the vegan package to address the high sparsity and compositional nature of the data structure. To evaluate the dissimilarities between samples with PE250 and PE300 data, principal component analysis (PCA) was performed on CLR transformed data. The statistical significance was assessed using the adonis2 function in the vegan package (999 permutations), based on Aitchison distance (i.e., Euclidean distance of the CLR-transformed data).

## Data and code availability

ITS2 raw sequencing data of HMP, GNHS, and MK-SpikeSeq were downloaded from NCBI (PRJNA356769), CNGBdb (CNP0002114), and ENA (PRJEB36435), respectively. All other source data, code, and analysis results are available on our <u>GitHub page</u>.

